# Too Soon to Save: structural uncertainty inverts the value of precautionary conservation action

**DOI:** 10.64898/2026.04.28.721297

**Authors:** Muhammad Umer Gurchani, José M. Montoya, Sacha Bourgeois-Gironde

**Affiliations:** Institut Jean Nicod (ENS, EHESS, CNRS), École Normale Supérieure, Paris; Centre National de la Recherche Scientifique (CNRS), Station d’Écologie Théorique et Expérimentale, Moulis; Université Paris 2 Panthéon-Assas, Paris; University of Haifa, Faculty of Law, Haifa

## Abstract

Conservation policy commonly assumes that acting early is inherently safer than waiting. Here, we show that this intuition can fail when ecological structure is uncertain and protection decisions are difficult to reverse. We compare a precautionary strategy that protects early under uncertainty with an adaptive strategy that learns before committing protection, across both synthetic and real ecosystems.

In synthetic ecosystems with uncertain trophic structure, the adaptive learn-then-commit strategy yields higher protected-area phylogenetic diversity than the main precautionary baseline (Rao’s *Q* PD = 5.23 versus 4.41, *P* < 10^−4^, Cohen’s *d* = 0.54) and higher functional diversity (1.39 versus 1.25, *P* < 10^−4^, *d* = 0.93), although it remains below the full-knowledge oracle (5.34 and 1.43, respectively). This adaptive advantage is greatest when errors in structural allocation are most costly, particularly in highly connected ecosystems. It is also stronger in highly modular systems, although this effect is secondary.

In a real ecosystem (North-East Atlantic fish communities), we find the same conditions for such an advantage: structural importance is largely decoupled from abundance (*ρ* = −0.05, *P* = 0.77), and trophic uncertainty declines markedly through time (*R*^2^ = 0.95, *P* < 10^−6^). Consistent with this mechanism, adaptive spatial allocation also outperforms a precautionary Marxan-like baseline in the empirical analysis (Shannon diversity 1.70 versus 1.44 at *K* = 10, *P* < 10^−5^).

Together, these results show that the value of waiting in conservation does not arise from delay itself, but from the opportunity to learn which components of an ecosystem matter most. When ecological structure is uncertain and protection is hard to reverse, precaution can lock conservation into avoidable mistakes.

## Introduction

The precautionary principle holds that lack of full scientific certainty should not be used as a reason for postponing measures to prevent biodiversity loss(1; 2). In practice, this often encourages front-loaded intervention: designate reserves early, commit scarce area quickly, and treat delay as inherently dangerous. That intuition is morally compelling, but it is not always strategically correct(7).

The hidden assumption behind early precaution is that protecting something now is always better than protecting later and better. That assumption is safe when conservation targets are effectively independent. It becomes much less safe when species are embedded in trophic networks, because the value of protecting one place depends on which interactions sustain the rest of the community(3). Under structural uncertainty, managers do not yet know which cells contain the species whose trophic roles matter most. If protection is also irreversible, acting early can permanently lock in the wrong spatial pattern.

This is an irreversibility-and-information problem in the sense of Arrow and Fisher(6): when better information is expected, preserving flexibility can have positive option value. Ecological applications have shown that learning can outperform immediate commitment when uncertainty is decision-relevant(8; 9; 14; 15). But existing arguments for delayed conservation rest primarily on growing budgets or restoration capacity(7). What remains largely unquantified is the cost of acting under *structural* ignorance about ecological interactions—a distinct mechanism, because structurally important species need not be the most abundant ones, so abundance-based protection can misallocate scarce effort even when budgets are fixed.

Here we ask under what ecological conditions early precautionary protection produces poorer long-run reserve outcomes than learning before commitment, and whether the structural prerequisites for such an adaptive advantage hold in a real marine system. We combine a partially observable decision framework for conservation planning in synthetic ecosystems with an empirical analysis based on ICES stomach-content networks and North Sea trawl data(28; 29; 18). We find that adaptive learning consistently outperforms early commitment, but the advantage is regime-dependent: it is greatest in highly connected and highly modular ecosystems, where structural misallocation is most costly, and smallest under strong disturbance, where rapid environmental turnover erodes the value of learning. Our central claim is that the value of waiting in conservation arises not from delay itself, but from the opportunity to learn which components of an ecosystem matter most—and that this opportunity is largest precisely in the structurally complex systems where precaution is most often invoked.

## Results

### Framework and ecosystem construction

We began with three synthetic ecosystems, each containing 8 species distributed across a 5 × 5 grid and linked by coupled trophic dynamics (Methods). To keep the construction biologically structured rather than arbitrary, we generated phylogenies under a Yule process(21), evolved traits under an Ornstein– Uhlenbeck process(22; 23), and translated trait differences into trophic interaction strengths using standard food-web logic(24; 25). The resulting ecosystems contained 5, 10, and 7 trophic edges.

The conservation problem is therefore partially observable. In every period, the true trophic matrix **A** remains hidden, and all implementable strategies receive only partial information about it: noisy abundance observations together with noisy evidence about predator–prey links. In that sense, all of the realistic strategies operate in the same underlying POMDP environment. What differs among them is not the uncertainty they face, but how they respond to it.

The adaptive strategy is the only one that uses this sequential structure directly. After each round of observations, it updates Bayesian posteriors over trophic edges and plans from its current belief state with POMCP(28). Because protection is irreversible and budget-limited, it does not simply decide where to protect, but also whether to protect now or preserve flexibility for later. Its reward weights are learned with maximum-entropy IRL(29) from a hindsight teacher, so the planner is trained to recover the trade-offs that would have mattered under the true interaction structure.

The two precautionary baselines, by contrast, do not use the problem in this sequential way. The naïve strategy uses only the initial noisy abundance snapshot, ranks cells by apparent abundance, and commits its full budget immediately. The Marxan-like strategy also commits immediately, but instead uses simulated annealing on the initial noisy data to choose a spatial set that improves early phylogenetic and functional complementarity. Neither baseline updates its view of trophic structure through time, and neither can revise its allocation after acting. Both therefore represent front-loaded precaution under structural ignorance.

The oracle plays a different role. It is not an implementable strategy, but a full-information upper bound. It knows the true trophic structure and uses trajectory-simulating simulated annealing to choose the best set of cells at *t* = 0. Its purpose is not to represent a realistic policy, but to show how far each implementable strategy remains from the best possible allocation under perfect knowledge.

Fig. 1 provides an overview of the problem studied here: conservation decisions are made under partial information about trophic structure, structural and abundance priorities can diverge across space, and the key strategic difference is between immediate commitment and learning before irreversible protection.

**Figure 1:**
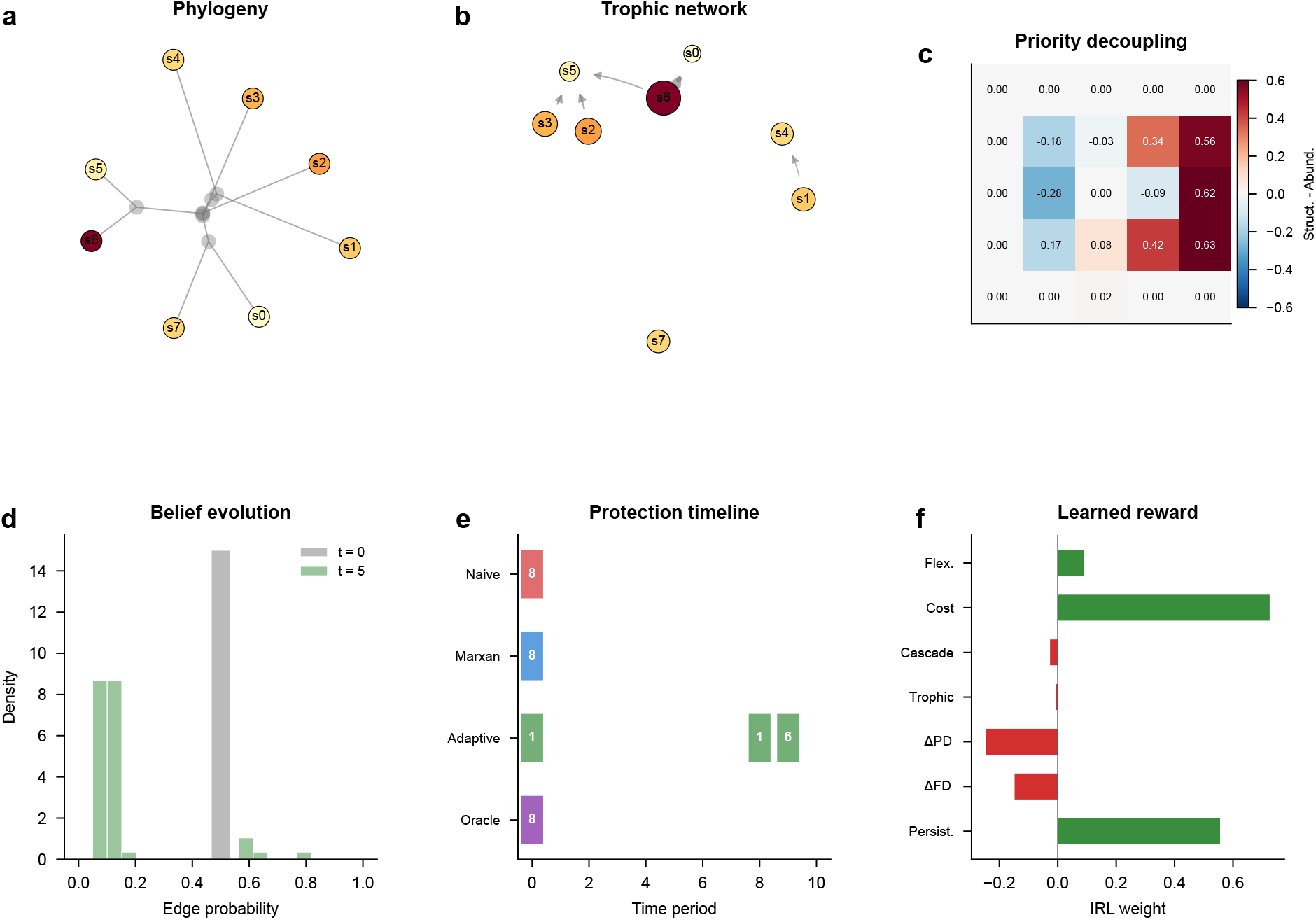
Framework for adaptive conservation under structural uncertainty. **a**, Yule phylogeny; tips coloured by keystone score (yellow = low, dark red = high). **b**, Trophic network; node size indicates keystone importance. **c**, Priority decoupling: cell-level difference between structural importance and abundance importance (red = structurally critical but low-abundance; blue = abundant but structurally replaceable). **d**, Belief evolution: edge-probability posteriors at *t* = 0 (grey) and *t* = 5 (green). **e**, Protection timelines across the four strategies. **f**, IRL-learned reward weights; cascade prevention dominates.

### Adaptive management outperforms precautionary strategies

Across the default synthetic experiments, the adaptive policy outperformed both implementable precautionary baselines on the two headline protected-area biodiversity metrics, while remaining below the full-information oracle (Fig. 2). Adaptive Rao’s *Q* phylogenetic diversity reached 5.23 [95% CI 4.78– 5.70], compared with 4.41 [3.99–4.86] for the Marxan-like baseline and 5.34 [4.86–5.85] for the oracle (*P* < 10^−4^, Cohen’s *d* = 0.54). Functional diversity (Rao’s *Q* on traits) showed the same ordering, with adaptive = 1.39 [1.35–1.42], Marxan-like = 1.25 [1.20–1.30], and oracle = 1.43 [1.38–1.47] (*P* < 10^−4^, *d* = 0.93). The naïve abundance-ranking strategy also performed well on phylogenetic diversity (5.01), because in these ecosystems structurally important species often coincide spatially with locally abundant species, but the adaptive policy retained a clear advantage on functional diversity. The gap between adaptive and oracle (approximately 2% of the oracle PD value) quantifies how much further gain would require full trophic knowledge rather than belief-state learning.

**Figure 2:**
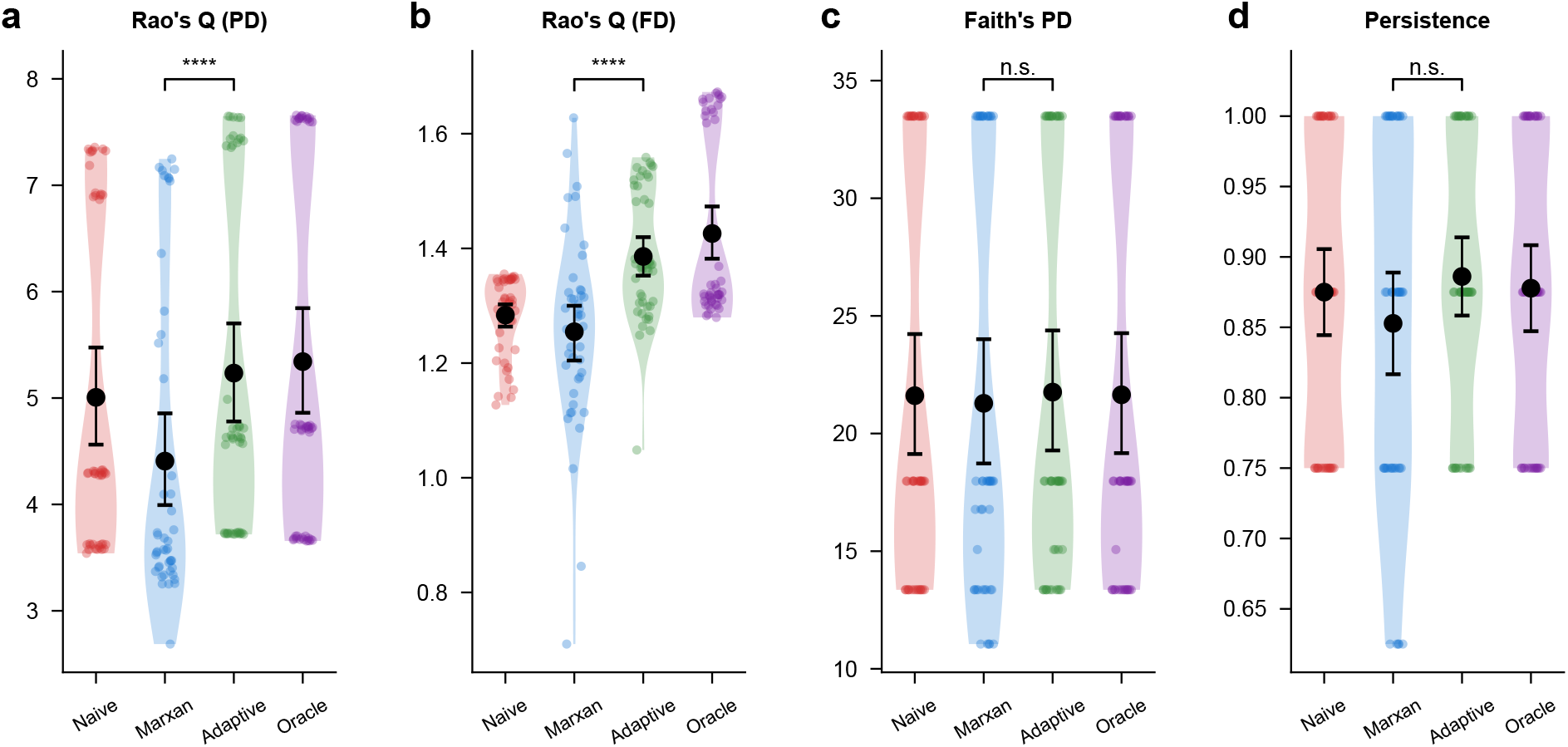
Adaptive management improves protected-area biodiversity outcomes. Violin plots of Rao’s *Q* phylogenetic diversity, Rao’s *Q* functional diversity, Faith’s PD, and protected-area persistence across the four strategies, with bootstrap means and 95% confidence intervals overlaid. The adaptive strategy exceeds the implementable early baselines on the two Rao’s *Q* metrics while remaining below the oracle upper bound.

### Adaptive gains are largest where structural mistakes are most costly

The size of the adaptive advantage varied markedly across ecological regimes (Fig. 3). The largest gains occurred in regimes where errors in spatial allocation were most consequential. In highly connected ecosystems, adaptive phylogenetic diversity reached 4.04 versus 3.12 for the Marxan-like baseline, a difference of ΔPD = +0.92 (*d* = 1.89, *P* = 0.0017). In highly modular systems, the adaptive advantage was the next largest (ΔPD = +0.60, *d* = 0.81), followed by the weak-keystone (+0.36), high-disturbance (+0.35), and low-disturbance (+0.27) regimes. In these settings, the ecological value of a cell depends strongly on the wider interaction structure, so protecting the wrong cells under uncertainty carries a larger long-run penalty.

**Figure 3:**
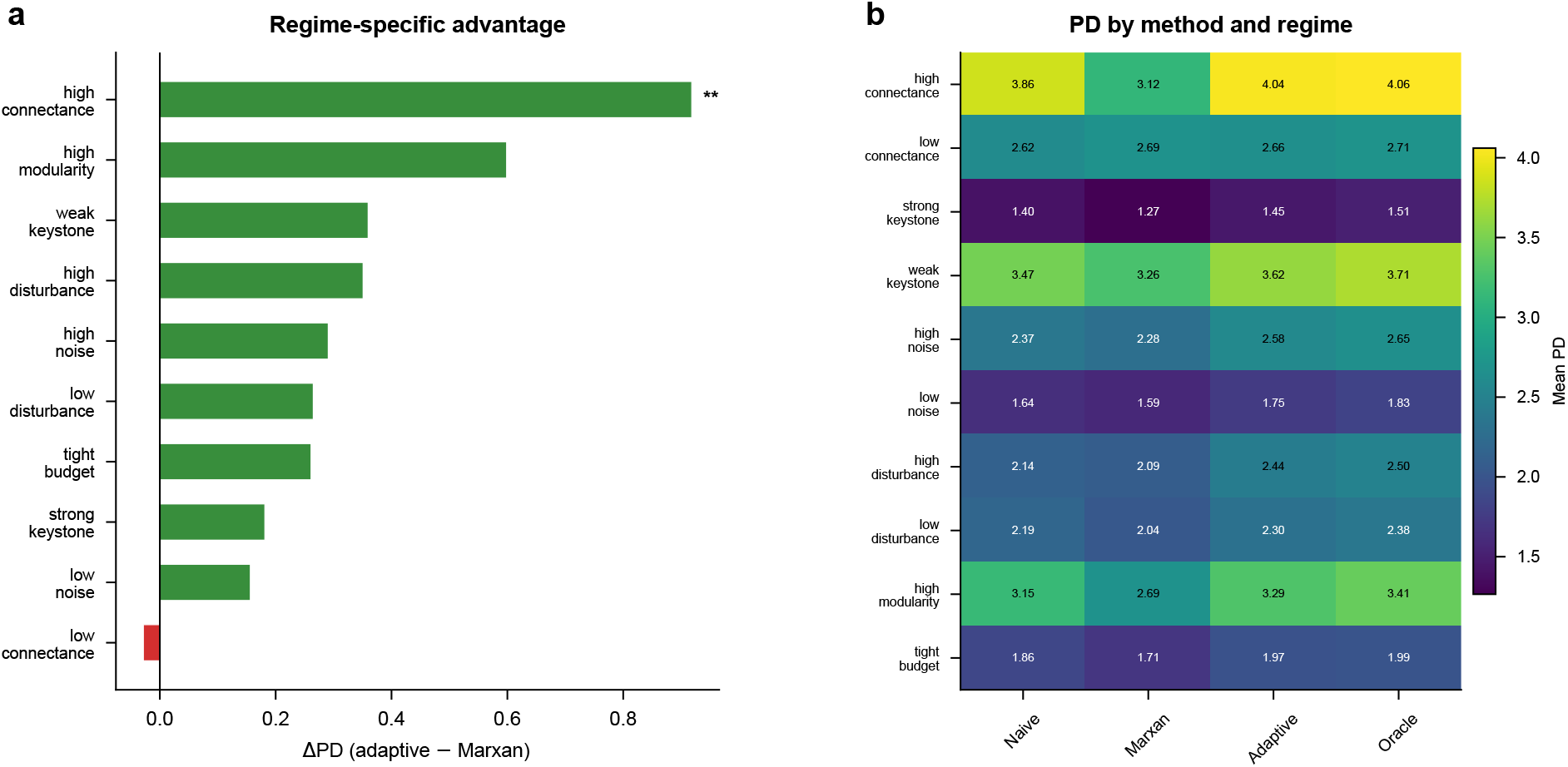
Adaptive advantage depends on ecological regime. **a**, Horizontal bars show ΔPD (adaptive − Marxan) across the ten regime modifications; asterisks mark regimes where the pairwise *P* -value crossed conventional thresholds. **b**, Mean protected-area Rao’s *Q* PD by method and regime. The adaptive gain is largest where trophic structure matters most for the allocation problem, especially in highly connected and highly modular systems.

Overall, the adaptive policy dominated in 9 of the 10 regimes, and the two precautionary baselines matched or narrowly beat it only in the low-connectance setting (ΔPD = −0.03). After Bonferroni correction across regimes, the high-connectance difference remained significant, while the remaining advantages were directionally consistent but noisier. The benefit of adaptive learning is therefore greatest not in every setting equally, but where structural uncertainty most strongly affects which cells ought to be protected: when networks are densely connected or modular, early commitment is most likely to lock in structurally misaligned reserves, and learning before committing recovers the largest share of the achievable gain.

### Empirical evidence for adaptive advantage

We then turned to the empirical analysis to ask whether the two conditions required by the synthetic model are present in a real marine system: first, that structural importance is not well captured by abundance alone; and second, that trophic structure becomes more informative through time. We tested both conditions using ICES stomach-content data for network reconstruction and independent trawl data for abundance, before comparing the four allocation strategies in a retrospective rectangle-allocation exercise.

Structural importance is not captured by abundance. We estimated species-level structural importance (SI) from probability-weighted posterior trophic networks inferred from stomach data and compared it with independent trawl CPUE. The association was essentially absent (Spearman *ρ* = −0.048, *P* = 0.77), indicating that abundant species were not, in general, the structurally most important ones (Fig. 4). This is the empirical analogue of the misalignment built into the simulations: abundance alone is a poor guide to the components of the system that matter most for trophic organisation. For clarity, the food-web visualization shows only the strongest posterior interactions, but the SI calculations use the full probability-weighted posterior network.

**Figure 4:**
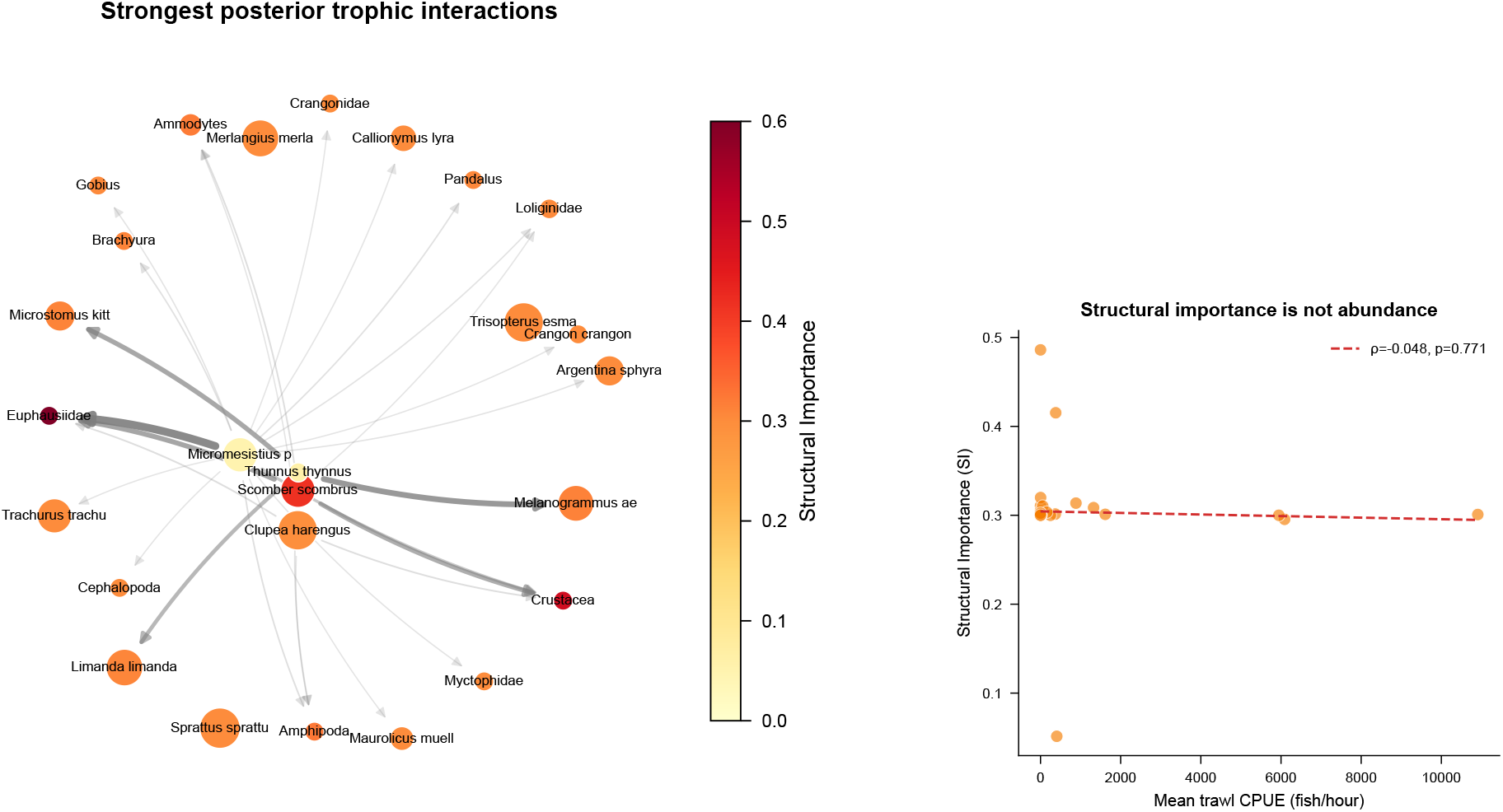
Empirical decoupling of structural importance and abundance. Left, strongest posterior trophic interactions among focal NE Atlantic taxa; edge width scales with posterior probability, node size with CPUE, and node colour with SI. Right, full species-level SI versus abundance comparison (*ρ* = −0.048, *P* = 0.77).

#### Trophic structure becomes more informative through time

Decade-by-decade Bayesian reconstruction showed a strong decline in mean edge entropy through time (*R*^2^ = 0.95, *P* < 10^−6^), meaning that uncertainty about predator–prey links narrowed as evidence accumulated. In parallel, SI rankings converged progressively toward the final-decade ordering, with Kendall’s *τ* increasing from 0.33 to 1.0 (Fig. 5). Together, these trends indicate that the system is learnable in the sense relevant to this paper: additional observation changes the inferred structural ranking of species, and does so in a progressively more stable way. Binary overlap with the final thresholded network remained sparse, but this is largely because the thresholded network itself is very small. In this empirical setting, the clearest signal of learnability lies not in recovering a dense binary graph, but in the reduction of uncertainty and the convergence of structural rank order.

**Figure 5:**
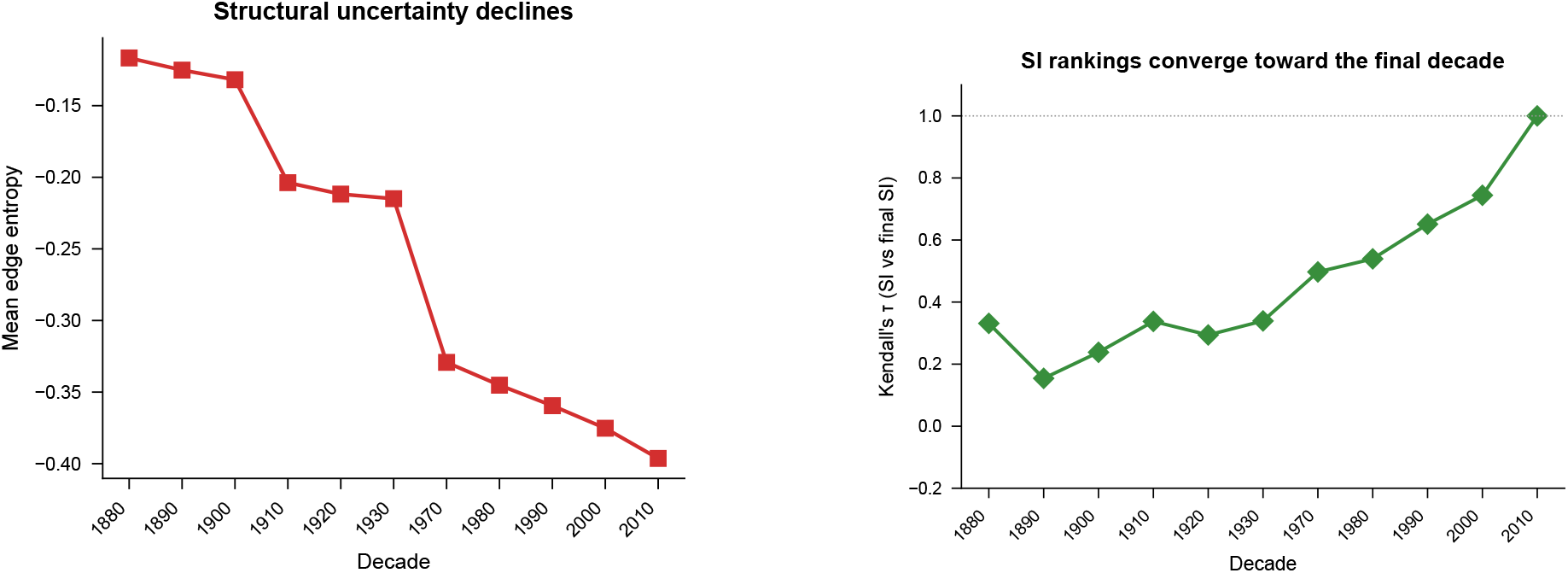
Trophic structure becomes progressively more informative through time. Left, mean posterior edge entropy declines across decades. Right, decade-specific SI rankings converge toward the final-decade ranking. These are the clearest empirical learnability signals in the sparse North-East Atlantic reconstruction.

#### Adaptive allocation also outperforms the implementable early baselines in the empirical analysis

Once the empirical oracle was defined as a true hindsight upper bound, the strategy comparison became much clearer (Fig. 6). At *K* = 10, the oracle achieved the highest Shannon diversity in the selected rectangles (*H*^′^ = 2.11), followed by the adaptive strategy (1.70), the Marxan-like baseline (1.44), and the naïve baseline (1.24). Thus, the adaptive strategy retained more diverse assemblages than either implementable early-commitment rule, while still remaining below the unattainable full-information benchmark. The adaptive advantage over Marxan was statistically strong (*P* < 10^−5^, Cohen’s *d* = 1.10), and the same ordering persisted across the full budget range. Importantly, this empirical ranking matched the simulation exactly (oracle > adaptive > Marxan > naïve; Spearman *ρ* = 1.0). The empirical message is therefore the same as in the synthetic analysis: when structural importance is decoupled from abundance and becomes clearer through time, learning before committing can improve spatial conservation outcomes.

**Figure 6:**
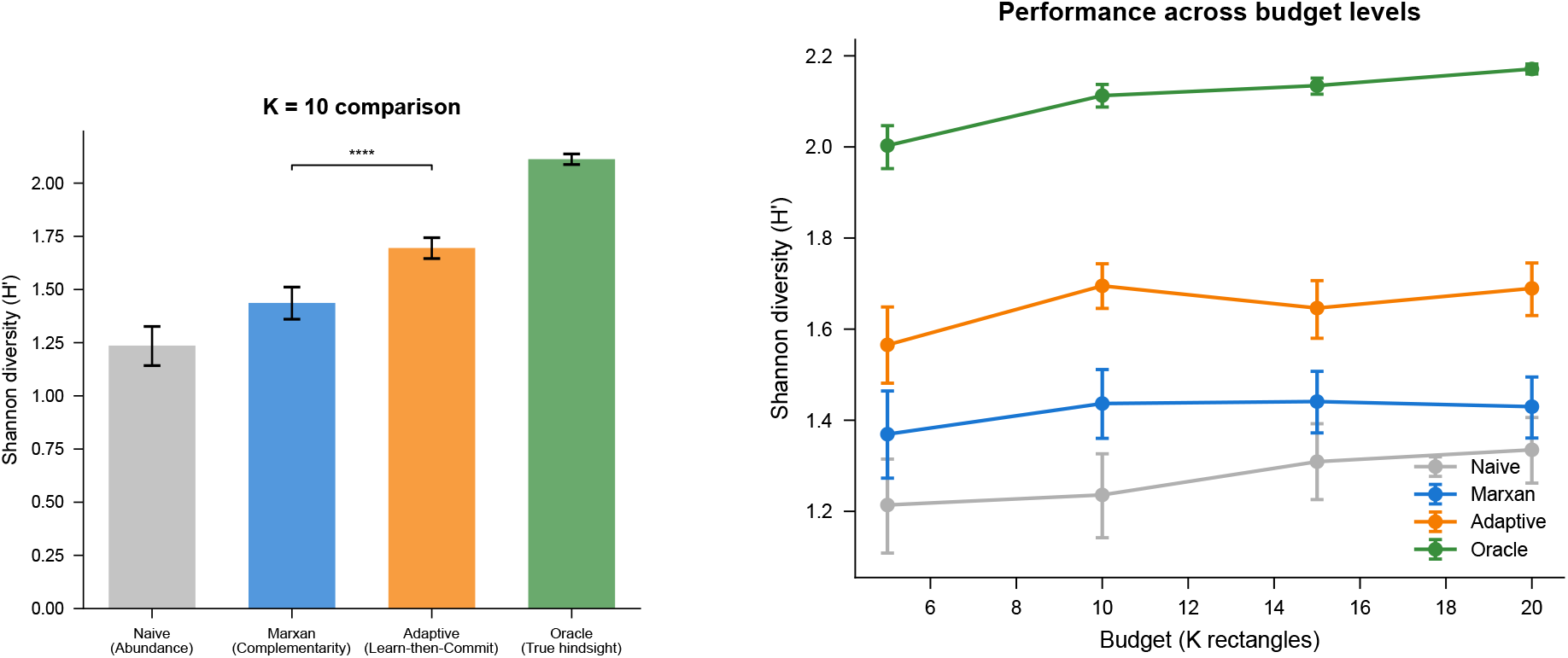
Empirical strategy comparison with a true hindsight oracle. Left, Shannon diversity by strategy at *K* = 10; the primary empirical policy comparison is adaptive versus Marxan. Right, performance across budget levels, showing a stable ordering of oracle > adaptive > Marxan > naïve.

### A secondary synthetic timing analysis

As a secondary analysis, we ran a synthetic factorial experiment that crossed five timing strategies with three budget levels. In that experiment, protection timing explained approximately 9.6× more variance in protected-area PD than total budget (*η*^2^ = 0.055 for timing versus 0.006 for budget), though neither factor reached conventional significance in this smaller design (*P* = 0.17 for timing, *P* = 0.71 for budget). Across budget levels, the best-performing fixed rule was the “late” schedule, which allocates protection after initial belief updating (mean PD = 4.21, 4.19 and 4.20 at budget 4, 8 and 12), while the endogenous POMCP policy remained close to this upper envelope (4.17, 4.19, 4.18). We interpret these numbers cautiously: they reinforce, but do not on their own establish, that irreversible commitment should be treated as a scheduling problem as well as a budgeting problem, and they are not one of the paper’s primary empirical claims.

## Discussion

Our central result is that precautionary conservation can be self-defeating under a specific, identifiable set of conditions. When trophic structure is uncertain and protection is irreversible, committing early does not merely forgo information—it permanently locks managers into spatial allocations that a better-informed decision would have avoided. The problem is not delay versus action. It is that “act now” and “act safely” can point in opposite directions when the ecological value of a site depends on interactions that are not yet known.

This finding modifies an existing theoretical tradition rather than contradicting it outright. Arrow and Fisher’s option-value argument (6) established that preserving flexibility has value when better information is expected and decisions are hard to reverse, and Iacona *et al*. (7) extended that logic to conservation triage. But prior formulations locate the value of waiting in growing budgets or future restoration capacity. Our mechanism is different: the value of waiting arises from *structural learning*—the progressive resolution of which ecological roles matter most—rather than from increasing resources. This distinction matters because it implies that even under fixed budgets, delay can improve outcomes, provided that the delay is used to reduce uncertainty about interaction structure. It also distinguishes our framing from the POMDP-for-ecology literature (14; 15), which treats partial observability primarily as a monitoring problem. Here, partial observability creates a *commitment* problem: irreversible protection under structural ignorance is costly not because the manager lacks data, but because the data that matter most—who eats whom, and which roles are replaceable—arrive after the decision window has closed.

The adaptive advantage is not universal, and its boundary conditions are informative. Three prerequisites must hold simultaneously. First, structural importance must be poorly predicted by abundance, so that simple observation-based rules misallocate protection. Second, learning must be fast enough relative to ecological turnover that the manager can resolve the relevant uncertainty before the system changes beyond recognition. Third, protection must be sufficiently irreversible that early mistakes persist rather than self-correct. When any one of these breaks down, the case for early commitment strengthens. In the synthetic experiments, the adaptive gain was largest in highly connected and highly modular systems, where a cell’s conservation value depends most strongly on the wider network, and essentially disappeared in the low-connectance regime, where the trophic graph is sparse enough that abundance alone is an adequate guide to structural importance. These are not isolated parameter sensitivities. They trace a coherent ecological logic: learning before acting matters most when the network makes mistakes expensive, and matters least when network structure is simple enough that early observation-based rules already approximate the right allocation.

The empirical analysis plays a different role from the simulations. It does not replicate the synthetic decision problem. Instead, it tests whether the *prerequisites* for adaptive advantage—decoupled structural importance and declining trophic uncertainty—hold in a real marine system. Both conditions are present in the North-East Atlantic data: structural importance was effectively unrelated to abundance (*ρ* = −0.05, *P* = 0.77), and posterior edge entropy declined strongly through time (*R*^2^ = 0.95). The retrospective allocation exercise then produces the same strategy ordering as the simulations (oracle > adaptive > Marxan > naïve). This convergence between synthetic and empirical analyses suggests that the mechanism we identify is not an artifact of the simulation design, but reflects a structural feature of food webs in which keystone roles are not telegraphed by abundance.

Several limitations define the scope of these conclusions rather than undermining them. The synthetic ecosystems are small (8 species, 25 cells) and the POMDP planner is far simpler than what operational conservation would require. The primary comparisons evaluate biodiversity within protected areas; ecosystem-wide outcomes may diverge if protection of a few structurally critical cells does not generate sufficient spillover to sustain unprotected populations at scale. The empirical analysis is retrospective, and the posterior trophic network is an estimate rather than ground truth. More broadly, our framework isolates trophic uncertainty from the many other dimensions of real conservation decisions— dispersal corridors, non-trophic mutualisms, governance constraints, and socio-economic trade-offs— each of which may interact with the timing question in ways we do not model. We therefore do not claim that waiting is generically better than acting. We claim that when the conditions we identify hold, the standard operationalisation of precaution—commit early, treat delay as risk—can be the less cautious choice.

These results suggest that conservation planners facing uncertain trophic structure should treat the timing of irreversible protection as a decision variable rather than a default. Systematic conservation planning (4) has framed reserve design primarily as a one-shot spatial optimisation. Our results reframe it as a sequential problem in which each action does two things: it protects biodiversity, and it determines whether future choices will be better or worse informed. In that framing, the relevant policy question is not whether to act, but whether acting now forecloses the opportunity to act better—and the answer depends on whether the ecological structure that determines “better” is still being learned.

## Methods

We used two linked analyses. The simulation analysis asks how a conservation manager should allocate irreversible protection while trophic structure is only partly known. The empirical analysis asks whether the same structural ingredients appear in a real marine system and whether a retrospective adaptive allocation rule outperforms simpler early-allocation baselines. The aim of the Methods is therefore not only to specify the code, but also to make clear how these two pieces fit together.

### Simulation decision problem

In the simulation, the manager repeatedly chooses whether to protect cells immediately or preserve budget for later decisions. Protection cannot be reversed, so waiting has value only if it leads to better targeting. Formally, the state is

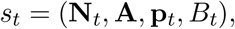

where 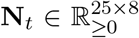 is the cell-by-species abundance matrix, **A** ∈ ℝ^8×8^ is the latent trophic interaction matrix, **p**_*t*_ ∈ {0, 1}^25^ is the irreversible protection mask, and *B*_*t*_ is the remaining budget. Actions are subsets of unprotected cells whose total cost does not exceed the remaining budget; the empty action is allowed and represents a decision to wait.

#### Ecological dynamics

We simulate multiplicative density-regulated trophic dynamics on a 5×5 grid. Protected cells receive a 40% boost relative to their baseline niche values, whereas unprotected cells are evaluated at 92% of baseline in each period. Growth combines density dependence, trophic interaction terms from the latent matrix **A**, nearest-neighbour dispersal (*m* = 0.01), Gaussian disturbance noise (*σ* = 0.08), and an explicit cascade penalty when the dominant keystone species declines in cells where it should otherwise persist. Abundances are truncated to [0.01, 50].

#### Observations and belief updates

The manager does not observe the trophic matrix directly. Instead, it receives noisy abundance observations and noisy evidence about predator–prey links. Abundance measurements are multiplicative Gaussian observations with period-specific noise *σ*_*t*_ = 0.3*/*(1 + 0.15*t*). Edge observations are Binomial counts with effort 20(1 + 0.1*t*) per predator–prey pair, with higher detection probability for true edges than for absent ones. The agent maintains Gaussian posteriors over abundances and Beta(1,1)-initialised posteriors over edge existence, and it tracks structural uncertainty from the corresponding posterior entropy and Beta moments.

#### Reward

The adaptive agent uses a linear reward,

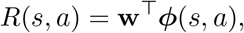

where ***ϕ*** contains seven interpretable features computed on the protected set: persistence, ΔFD, ΔPD, trophic importance, cascade prevention, protection cost, and flexibility loss. Weights are learned rather than hand-set, so the model can recover which trade-offs matter most for hindsight-optimal conservation.

### Planning, teacher policy and oracle benchmark

Given its current belief state, the adaptive agent plans with Monte Carlo belief-tree search (POMCP)(28). The planner uses 120 simulations per decision, search depth 3, UCB1 exploration constant *c* = 1.5, discount *γ* = 0.95, and 30 belief particles.

To obtain an interpretable reward, we train the adaptive policy with maximum-entropy IRL(29) on demonstrations from a hindsight teacher. The teacher re-evaluates earlier states using the true trophic structure and compares its chosen action with six sampled alternatives per state during the first two exploration periods. The oracle benchmark is stricter still: it uses the true trophic matrix and trajectory-simulating simulated annealing (10 restarts, 3000 iterations) to choose the full protected set, which is then committed at *t* = 0. In other words, the teacher provides training examples, whereas the oracle provides the upper bound.

### Synthetic ecosystems and simulation experiments

Each synthetic ecosystem contains 8 species on a 5 × 5 grid. Phylogenies are generated under a Yule pure-birth process(21) with *λ* = 1.0, and four continuous traits evolve under an Ornstein–Uhlenbeck process(22; 23) with *α* = 0.6 and *σ* = 0.7. Trait differences are then converted into directed trophic interactions(24; 25).

Species are placed in the grid naturally, without any post-hoc manipulation of keystone habitat: a species’ spatial distribution is determined by its evolved trait values through the Gaussian niche model, and its keystone score is determined by its position in the trophic network. Cell-level structural importance is the normalised sum of positive keystone scores among species present in a cell, whereas abundance importance is the normalised total initial abundance in that cell. In the three default ecosystems, the cell-level correlation between structural and abundance importance was 0.77–0.91, so that abundance-based ranking captures part of the structural signal but not all of it: any decoupling between the two quantities arises endogenously from the ecological model rather than from forcing, and is weaker than in a deliberately misaligned setup.

Biodiversity outcomes are evaluated on the protected set using Rao’s *Q* for phylogenetic diversity and functional diversity(26; 27), together with supplementary coverage metrics such as Faith’s PD, persistence and biomass. The default synthetic comparison uses three ecosystems with 15 replicates each. A separate factorial analysis varies timing strategy and total budget; we report that timing analysis as a secondary synthetic result rather than as one of the paper’s main empirical claims.

### Empirical data and network reconstruction

The empirical analysis combines two independent data sources. Trophic interactions come from the ICES stomach-content database(18), whereas abundance and biodiversity evaluation come from NS-IBTS trawl surveys aggregated to ICES rectangles. Species are harmonised through WoRMS AphiaID identifiers. The focal empirical set is the intersection of species that appear in the stomach records (as predator or prey) and in the trawl data. Stomach data are used only for network reconstruction; trawl CPUE is used only for abundance mapping and retrospective evaluation.

For each decade and each predator species, we update predator–prey links with a Beta-Binomial model. If a predator has *n* stomach observations in a decade and prey *j* appears *s* times, the corresponding edge posterior is updated by adding *s* successes and *n* − *s* failures to a Beta(1,1) prior. This yields decade-specific posterior edge probabilities and a transparent measure of how structural uncertainty changes through time.

The main empirical structural-importance calculations use the full probability-weighted posterior network, not only a hard threshold. Binary thresholded networks are retained only for diagnostic comparisons such as Jaccard overlap and the exploratory connectance sweep. For visual clarity, the food-web figure in the main text shows the strongest observed posterior interactions rather than all possible links.

### Empirical retrospective allocation analysis

To mirror the simulation, we compare four rectangle-allocation rules. The naïve strategy chooses earliest-decade rectangles by abundance alone. The Marxan-like strategy uses simulated annealing on earliest-decade species complementarity. The adaptive strategy chooses a decision decade from the entropy trajectory and then optimizes rectangles with decade-appropriate SI from the corresponding posterior network. The empirical oracle is a true hindsight upper bound: it chooses earliest-decision rectangles to maximize realized final-decade Shannon diversity by simulated annealing over the retrospective evaluation objective.

Protected-rectangle performance is evaluated in the final trawl decade with species richness, Shannon diversity, persistence and total CPUE. In practice, Shannon diversity is the most informative empirical outcome because richness is saturated across protected rectangles in this dataset.

### Statistics

Pairwise strategy comparisons use Mann–Whitney *U* tests with Bonferroni correction. Effect sizes are reported as Cohen’s *d*, and confidence intervals are 95% bootstrap intervals based on 2000 resamples. Timing–budget diagnostics are summarized with two-way ANOVA effect sizes (*η*^2^). In the empirical notebook, the timing and connectance diagnostics are treated explicitly as exploratory rather than confirmatory.

## Acknowledgements

This research was co-funded by the European Union (GA#101059915 – BIOcean5D). Views and opinions expressed are those of the authors only and do not necessarily reflect those of the European Union or the granting authority.

## Author contributions

M.U.G. conceived the study, developed the POMDP–IRL framework, implemented all simulations and analyses, and wrote the manuscript. J.M.M. provided ecological expertise, guided the empirical validation, and edited the manuscript. S.B.-G. contributed the decision-theoretic framing and edited the manuscript.

## Competing interests

The authors declare no competing interests.

## Data and code availability

ICES stomach data: https://ices.dk/data/data-portals/Pages/stomach.aspx. NS-IBTS trawl data: https://www.ices.dk/data/data-portals/Pages/DATRAS.aspx. All code used to generate the results and figures in this study is available at https://github.com/Gurchani/adaptive-conservation-pomdp under an MIT license.

## Notes

### Competing Interest Statement

The authors have declared no competing interest.

